# An Abundant New Saponin-Polybromophenol Antibiotic (CU1) from *Cassia fistula* Bark Against Multi-Drug Resistant Bacteria Targeting RNA Polymerase

**DOI:** 10.1101/2020.11.04.369058

**Authors:** Asit Kumar Chakraborty, Sourajit Saha, Kousik Poria, Tanmoy Samanta, Sudhanshu Gautam, Jayanta Mukhopadhyay

## Abstract

Spread of multidrug-resistant infections is a threat to human race and need for new drug development is great. Bark ethanol extract of *Cassia fistula* inhibited MDR bacteria isolated from Ganga River water, human and animal. On TLC, a gray colour major band ran fast (CU1; 6.6% of bark and ~30% of crude extract) which quickly purified on HPLC C_18_ column at 3 min. Chemical assays suggested a triterpene linked to polyphenol known as saponin. CHN Elements analysis (35.9% C; 5.5% H) did not identified nitrogen suggesting a polyphenol or glycoside. VU-Vis spectra gave high peak at below 200nm with a secondary peak at 275nm with minor hinge at 578nm indicating a fused ring with bromo-polyphenol. CU1 Mass (897 Daltons) with fragments of 515, 325, 269, 180 daltons and six halogen substitutions reflected by 82 molecular mass of DBr deviate six larger fragments. FT-IR suggested broad band at 3500-3000 cm^−1^ for −OH where as two strong peaks at 1552cm^−1^ for aromatic C=C and 1408cm^−1^ for phenol. Proton-NMR confirmed polymeric phenol at δ 4.86-4.91 ppm and tetratet at δ3.57-3.618 ppm with phenolic bromo-substituents. Carbon-NMR identified a strong peak at δ=23.7ppm for many C-Br and at 165ppm for a polybenzoid compound. CU1 inhibited the RNA Polymerase of *E. coli* (IC_50_=23μg/ml) and *M. tuberculosis* (IC_50_=34μg/ml) but not DNA polymerase. Gel shift assays demonstrated that CU1 drug interacted with enzyme and inhibited its binding to open promoter complex. Thus, CU1 phyto-chemical is an alternative safer and low cost drug against MDR-TB as well as other MDR pathogens.

## Introduction

Multidrug-resistant pathogenesis has caused a serious threat to society where anyone is susceptible to AMR (antimicrobial resistance) disease if immuno-system is weak. AMR disease is potential threat to life as most common antibiotics (ampicillin, streptomycin, chloramphenicol, ciprofloxacin, cefotaxime, erythromycin etc) did not able to kill bacteria due to presence of *mdr* genes (*bla, amp, cat, acc, aad, str, aph, sul, mcr, erm, cfr, dhfr*) in plasmids followed by chromosomal mutations (gyrAB, porB, L4, and rRNA) [1–3]. MDR genes *bla* homolouges (TEM, SHV, CTX-M, OXA, VIM, CMY) degrades ampicillin, cefotaxime and meropenem differentially but blaNDM1 enzyme degrade all, whereas CAT enzyme acetylates chloramphenicol only but AAC acetylating enzymes are heterogeneous and inactivate many other drugs. APH enzymes phosphorylate many drugs like streptomycin, azithromycin and amikacin to make inactive drug derivatives. Few dozens *mdr* genes were isolated and sequenced but million mutated isolates with potential higher drug inactivating capabilities were documented in human infections caused by *Salmonella, Pseudomonas, Acinetobacter, Klebsiella, Bacillus* and *Escherichia* species [4–7]. Importantly, small R-plasmids with 1-3 *mdr* genes have combined with F’-conjugative plasmid (62.5kb) giving more space to multiple (5-15) *mdr* genes as well as drug-efflux genes and many recombinases and IS-elements. Such bacteria can donate the *mdr* genes to other environmental bacteria by conjugation where we found 45% ampicillin resistance bacteria in Ganga River water and Digha sea water but reported clinical isolates were >95% ampicillin resistant [8]. Obviously question arises why such *mdr* genes creation in bacteria? Scientists predicted that gut bacteria were in symbiosis with human intestinal cells that would be die unless bacteria could supply vitamins. Vitamins as coenzymes (NAD, FAD, Biotin, PALPO, THFA) were component of ten thousands enzymes who made bio-molecules (amino acids, nucleotides, fatty acids and sugars) in our body [9]. Other words, intake of huge oral antibiotics since 1928 killed the gut bacteria and might be a signalling from intestinal cells secreting ILs and CKs (interleukins and cytokines) had helped bacteria to rearranged its plasmids and transposons to create *mdr* genes so that gut bacteria could live even we used to take multiple doses of oral antibiotics during fever and cough. Due to overwhelming drug void, drug companies are now not investing in new drug discovery against bacteria because it appears the half life for new drug may be few months and then a new *mar* gene will appear to destroy the drug. For new drug discovery it took two billion dollar and investors are in confusion and fear how they will recover the money from market! In such scenario, G-20 Leaders and WHO have approved a new dimension in drug discovery against multidrug-resistant bacteria from household medicinal plants [10–12].

*Cassia fistula* (Fabaceae family) is a deciduous medium sized tree (10-20m hight and 1-2m girth) cultivated in India for medicinal and decorative purposes and is also common throughout the Gangetic vally and sub-Himalayan tracts including other Asian countries like Srilanka, Thiland and Nepal [13]. The different parts (flower, root, bean, ripe fruit pulp, seed and leaves) are used in curing variety of infections and ailments [14]. A 3.5KD small protease inhibitor (Fistulin) was purified from leaves of *Cassia fistula* having anti-microbial activities [15]. Many chemical constituents have been analyzed from *Cassia* bean and tree parts [16–21]. We first reported the role of *Cassia fistula bark e*xtract in inhibiting superbugs [12]. In this communication, we have extended such studies purifying the active chemical and identifying its target as RNA polymerase.

In bacteria, a single RNA polymerase could transcribe all rRNA, mRNA and tRNAs whereas in eukaryotes three RNA polymerases like Pol I, Pol II and Pol III transcribed those RNAs respectively and also another mitochondrial RNA polymerase was characterized [22–24]. Bacterial RNA polymerase was inhibited by anti-tuberculosis drug refampicin which was useless in many MDR-TB cases now. Eukaryotic Pol-II was selectively inhibited by α-amanitin whereas Pol-I was insensitive to α-amanitin and Pol-III was moderately inhibited at high dose (50μg/ml). RNA polymerases were purified by DEAE-Sephadex column chromatography and were very labile to heat and EDTA. Presently, affinity column was used to quickly purify the enzyme and stored in 50% glycerol [23]. α-^32^P −UTP or H^3-^UTP was used to assay DNA template-dependent incorporation of nucleotide as RNA where RNAase treatment abolished the activity but not DNase treatment. However, specific RNA binding riboprobe binds the RNA produced and DNase treatment to remove DNA template and fluorescence detection method correctly detects the enzymatic activity in routine RNA polymerase ribo-probe assay [25]. We are influenced by herbal-drugs of ancient Indian Hindu Civilization described in Sanskrit books Charaka Samhita, Sasruta Samhita and Atharva Veda. We described here purification and characterization of phyto-chemical from *Cassia fistula* bark that interact with RNA polymerase inhibiting open-complex formation and RNA synthesis.

## Material and Methods

### Collection of bark and chemical extraction

We isolated bark from a medium sized tree (8m) and bark was thick (8mm). We cut 5 mm × 5 mm sized pieces of bark with the help of a knife (zatei) as motorized grinding generated heat to destroy active phyto-chemicals. The extraction was done with ethanol (1:5; w/v). Overnight extraction was good but 2-3 days extraction was found helpful. Ethanol extraction was good but methanol extraction yielded more CU1 than CU3 phytochemicals. Dried bark is good but fresh bark was also not harmful but some water (10%) soluble chemicals were contaminated. Dried extract was active for months in the refrigerator but ethanol extract at room temperature (35°C) was inactivated quickly. We recovered ~11g solid chemicals from 200g bark in 1000ml methanol extraction and evaporation [26].

### Preparative Thin Layer Chromatography

We used 20×15cm eight Silica gel plates for purification using 100ml solvent (methanol:water:acetic acid; 50:40:10) in glass jar with cover lid. It was found that CU1 chemical ran fast near solvent front and get clustered with lines with slow moving due to association of chemicals. From stock dry mass, we dissolved 1g solid in 10ml ethanol, clarified by 10000rpm for 5min and loaded onto silica plate (1cm wide). The ascending chromatography was done for 65min, dried the plates at room temperature, scratched the gray areas and suspended in 20ml ethanol and dissolved chemical was ran another TLC. Scraped the dark chemical again, and ultimately dried at room temperature (60mg CU1 recovered per preparation). Similarly, CU2 and CU3 phyto-chemicals were purified under UV-shadow. About 95-99% pure CU1 was obtained by repeated TLC [8].

### High Performance Liquid Chromatography

5mg TLC-purified active sample dissolved in 0.5ml methanol, filtered through a membrane filter and 0.1ml sample was loaded onto a HPLC C-18 column equilibrated with methanol 0.5ml fractions were collected and major active peak (retention time-3min) was collected and vacuum dried [29].

### Phyto-chemicals assays

Crude concentrated ethanol extract or TLC-purified chemicals were used for biochemical assays. TLC-purified CU1 chemical dissolved in water before assays and if ethanol extract used then added water during assay to lower ethanol concentration. *Assay of Saponins*: 0.5 ml plant extract + 2ml methanol + 2ml water. Formation of persistent foam on the surface was taken an indication for saponin. *Assay of Tannins*: 2ml bromine water + 0.3 ml plant extract. Decolouration indicates tannins, *Assay of Flavones*: Alkaline Reagent Test: 2◻ml of 2% NaOH solution was mixed with 0.4ml plant extract. A concentrated yellow colour was produced, which became colourless when added 2 drops of diluted HCl to mixture. *Test for glycosides*: Liebermann’s Test: Added 2 ml of acetic acid and 2◻ml of chloroform with plant extract. The mixture was then cooled and added H_2_SO_4_ concentrated. Green colour showed the entity of aglycone and glycosides. *Test for Terpenoids*: 0.5ml chloroform + 0.5 ml sulfuric acid concentrated + 0,5 ml plant extract + 0.5 ml water. Boiled the mixture. A grey colour indicates triterpenes. *Test for steroids*: 0.4ml chloroform + 1ml plant extract + few drop concentrated sulphuric acid. Red colour indicates steroids. *Test for alkaloids*: Alcohol extracts were diluted with dilute HCl (1:5v/v) and treated with few drops of Mayer`s reagent. A reddish brown or orange precipitates indicates the presence of alkaloids [27,28].

### Fourier Transformed Infra Red Spectroscopy

5mg HPLC-purified dry active chemical was mixed with 200mg IR-grade KBr and the tablet was prepared at 13mm Die SET (Kimaya Engineers) at 10Kg/cm^2^. Spectra were taken with a Perkin Elmer Spectrum 100 FT-IR Spectrometer (Serial no. 80944) for 10 min [30].

### NMR Spectrometry

Proton-NMR spectrometry was performed in CDCl_3_, D_2_O and CD_3_OD (4mg/ml CU1) for 10min. Carbon-NMR was performed in D_2_O only (20mg/ml CU1) in 500MHz JNM-ECX500 Spectrometer.

### Assay for anti-bacterial activities

We routinely used *E. coli-*KT-1_mdr and *P. aeruginosa*_PB-2_mdr for assay. The bacteria were resistant to ampicillin, ciprofloxacin, sulfamethoxazole, linezolid, vancomycin and moderate resistant to chloramphenicol, cefotaxime and streptomycin but sensitive to tigecycline and meropenem [8]. We also used linezolid, meropenem and amikacin resistant bacteria isolated from Ganga River water and Bay of Bengal sea water.100μl over night culture of MDR-bacteria was spread onto 10cm LB+ 1.5% Agar plate and 6mm holes were done and 40μl ethanolic solution of phyto-chemical added. In blank we loaded 40μl ethanol which usually did not interfere assay but rarely few strains of MDR bacteria gave minor lysis zone which likely was recovered (re-growth) during extended assay period. We determined >15mm lysis zone was good anti-bacterial activity where 20-30mm lysis zone could be detected for standard 20 μl of chloramphenicol (34mg/ml) or streptomycin (50mg/ml) or meropenem (1mg/ml) [12].

### Assay of Escherichia coli RNA Polymerase using ^3^H-UTP

The assay used to measure RNAP activity was performed as described by Lowe et al (1979) with minor modification [32]. The reaction (Rx 40μl) was performed with transcription buffer (40 mM Tris-HCl pH 7.9, 200 mM NaCl, 10 mM MgCl_2_, 0.1mM EDTA, 14 mM β-ME, 200nM each ATP, GTP and CTP. 50μM UTP, 2μCi ^3^H-UTP (BRIT, Hyderabad, India), 1.5μg calf thymus DNA and 1U RNA polymerase at 37°C for 20 min. The reaction mixture was spotted onto DEAE-paper pre-socked with 5mM EDTA. The filter paper washed with 5% di-sodium hydrogen phosphate, thrice with water and finally with 95% ethanol and dried. The filters were placed into scintillation vials containing 10 ml toluene-based scintillation fluid and counts were recorded on a Tri-CARB 2900TR scintillation counter. Data was normalized and plotted. Refampicin (10mg/ml in basic ethanol) was used as standard drug and 11 phyto-chemicals (4 times TLC purified; ~10mg/ml in DMSO) were used as experimental.

### Mycobacterium RNA polymerase purification and the bacterial strains used

*E. coli* strains BL21 (DE3) and C43 (DE3) have been used for the purification of *Mtb* RNA polymerase (RNAP) σ^A^holo and *Mtb.*σ^A^ respectively. For the purification of these proteins, respective *E. coli* cells have been transformed with their respective plasmids and were plated. The colonies obtained were then cultured in 2xYT medium (16g Tryptone, 10g Yeast Extract, 5g NaCl, pH 7.0) in presence of suitable antibiotics. The cultured cells were then pelleted down at 5000 rpm in a Sorvall cold centrifuge and lysed to purify the over expressed proteins of interest using specific purification protocols. *In vivo* assembled *Mtb* RNAP holo enzyme was purified as in Banerjee *et al.,* 2014 as also purification of recombinant σ^A^:Recombinant σ^A^ [33].

### α^32p^-UTP based in vitro transcription assay

Radioactivity based in vitro transcription assay was done as in Banerjee *et al.,* 2014. To perform this assay 300 nM *Mtb.*RNAP σ*A* holo was incubated in 1200 nM σ*A* in 2μl transcription buffer (45 mM Tris-HCl (pH8.0), 70 mM KCl, 1 mM DTT, 10% Glycerol, 5 mM MgCl_2_ and 1.5 mM MnCl_2_) at 25**°** Celsius for 15 minutes. 200nM DNA template (containing *sinP3* promoter) was added to the reactions along with 0, 10, 20, 40, 80, 160 μg/ml of the phyto-chemical CU1, and incubated at 37° Celsius for 15 minutes to form open promoter complex (RP_o_). Heparin (0.25mg/ml) was added to the reactions to inhibit non-specific RNAP-DNA complex. RNA synthesis was initiated by addition of NTP mix (final concentration 250μM of ATP, GTP and CTP and 10 μM UTP containing 5μCi α^32p^-UTP) and incubated for elongation at 37**°** Celsius for 30 minutes. The reactions were terminated by addition of 2μl FLB dye (80% formamide, 10 mM EDTA, 0.01% Bromophenol Blue, 0.01% Xylene Cyanol). Products were heated at 95° Celsius for 5 min, chilled on ice, and resolved by 12% Urea-PAGE, then scanned by storage phosphor scanner (Typhoon trio+, GE Healthcare) and quantified using Image Quant TL software.

### Fluorescence based in vitro transcription assay

Fluorescence based in vitro transcription assay was done as in Datta *et al.*, 2016 [34]. 400nM *Mtb.* RNAP*σ^A^*holo was incubated in 2μl transcription buffer (50 mM Tris-HCl (pH 8), 100 mM KCl, 10 mM MgCl_2_, 1mM DTT, 50nM BSA and 5% glycerol) at 25**°** Celsius for 15 minutes to reactivate. 5ng/μl of pUC19-*sinP3* plasmid DNA along with 0, 10, 20, 40, 80 μg/ml of the phytochemical CU1, was further added and incubatedaa t 37° Celsius for 20 min to form open promoter complex (RP_o_). RNA synthesis was initiated by the addition of NTP mix (final concentration 250μM of ATP, GTP, CTP and UTP) and incubated for elongation **at** 37°Celsius for 30 minutes. The reaction was stopped by adding 5U of RNase-free DNase I (Thermo Scientific), followed by incubation for 10 min at 37° Celsius. RiboGreen dye (Invitrogen, Carlsbad, CA) diluted to 200 folds in TE buffer (10 mM Tris-HCl (pH 8.5), 1 mM EDTA) was prepared. After DNase digestion, the samples were incubated for 5 min at 25**°** Celsius with the prepared RiboGreen dye solution in TE buffer, so that the reaction mix get 10-fold diluted. Fluorescence intensities of the samples were monitored using a spectrofluorometer (Photon Technology International, HORIBA Scientific, Edison, NJ) at excitation and emission wavelengths of 500 and 525 nm, respectively [25].

### Electrophoretic Mobility Shift Assay

Electrophoretic Mobility Shift Assay (EMSA) was done as in Rudra *et al.,* 2015 [35]. Cy5 labelled primer for the *sinP3* DNA fragment was used for this assay. The promoter DNA fragment was amplified using PCR and the Cy5 labelled amplified DNA was precipitated with an equal volume of isopropanol and 0.1 volume of 3M Na-acetate (pH 5.3). For the assay, 1.2μM *Mtb.* RNAP was incubated at 25**°** Celsius in 2μl transcription buffer (45 mM Tris-HCl (pH8.0), 70 mM KCl, 1 mM DTT, 10% Glycerol, 5 mM MgCl_2_ and 1.5 mM MnCl_2_) for 15 minutes to reactivate. 400nM Cy5 labelled *sinP3* promoter containing DNA fragment and heparin at 0.25mg/ml was then added to reactions along with 0, 5, 10, 20, 40, 80 and 160 μg/ml of the phyto-chemical CU1 and incubated at 37° Celsius for 30 minutes to form open promoter complex (RP_o_). The reactions were then resolved in 5% native PAGE in 1xTBE buffer (89 mM Tris base, 89 mM Boric Acid, 2 mM EDTA) for 90 minutes at 4° Celsius and then scanned and examined **by** Typhoon trio+ (GE Healthcare) scanner using Cy5 channel.

### In vitro replication assay

In vitro replication assay was done as in Rudra *et al.,* 2015 [35]. Single-stranded DNA template *pUC19-*sinP3 and its complementary Cy5 labelled primer were annealed in annealing buffer (50 mM Tris.HCl pH 7.5 and 100 mM NaCl) by heating to 95*°* Celsius followed by gradually cooling down to 25**°** Celsius. The sequence of the Cy5-sinP3 forward primer is: 5’-/Cy5/CAG CCA GAA GTC ATA CCG-3’. 50 nM annealed DNA template was incubated with 0, 10, 20, 40, 80 and 160 μg/ml of the phyto-chemical CU1 in respective reactions, at 37° Celsius for 5 minutes. 0.2 units of Klenow fragment (NEB) and 0.25mM dNTP mix were added to each reaction and were incubated at 37° Celsius for 1 minute. The reactions were terminated by addition of 2μl FLB dye (80% formamide, 10 mM EDTA) and heated at 95° Celsius for 2 min then chilled on ice. Products were resolved by 12% Urea-PAGE and the gel was scanned in Typhoon trio+ (GE Healthcare) scanner using Cy5 channel.

## RESULTS

### Anti-bacterial activities of CU1 and other phyto-chemicals from *Cassia fistula* bark

Anti-bacterial activity was previously reported for ethanol extract of *Cassia fistula* bark [8,12]. CU1 and CU3 anti-bacterial activities of TLC-purified phyto-chemicals were comparable but CU2 is less active. Five time concentrated extract or dried solubilised extract (10mg/ml in DMSO) showed antibacterial activity with 15-20mm lysis zone during Kirby-Bauer assay as compared to 20μl of 34mg/ml chloramphenicol or 20μl of 1mg/ml meropenem (20-25mm) using *Escherichia coli* KT-1_mdr (accession no. KY769881) or *Pseudomonas aeruginosa* DB-1_mdr (accession no. KY769875) (figure-2). We searched local area of Midnapore (22.658 Lattitude × 87.593 Longitude) and selected *Cassia fistula* bark (phyto-chemical named CU), *Suregada multiflora* (local name *Narenga*; phyto-chemical named NU), *Jatropha gossypiifolia* (local name *Varenda*; phytochemical named VU) as valuable source for new drugs to cure superbug infection where no antibiotics work. Indian spices like labanga (*Syzygium aromaticum* flower bud; named LU) and Derchini (*Cinnamomum zeynalium* bark; phyto-chemical named DU). During TLC purification, CU1 runs fastest moving chemical and CU2 is slower than CU1 and CU3 is slower than CU2 and much slower than CU1. We termed heterogeneous phytochemicals to explain the different derivatives of an extract or combination of 2-3 phyto-extracts from different plants and such extracts are good anti-bacterial drug and non-toxic [8]. Due to the multi-component nature of chemical structures, such heterogeneous phyto-antibiotics hard to become drug-resistant generating of *mdr* genes as well as activation drug-efflux genes.

**Fig.1.**
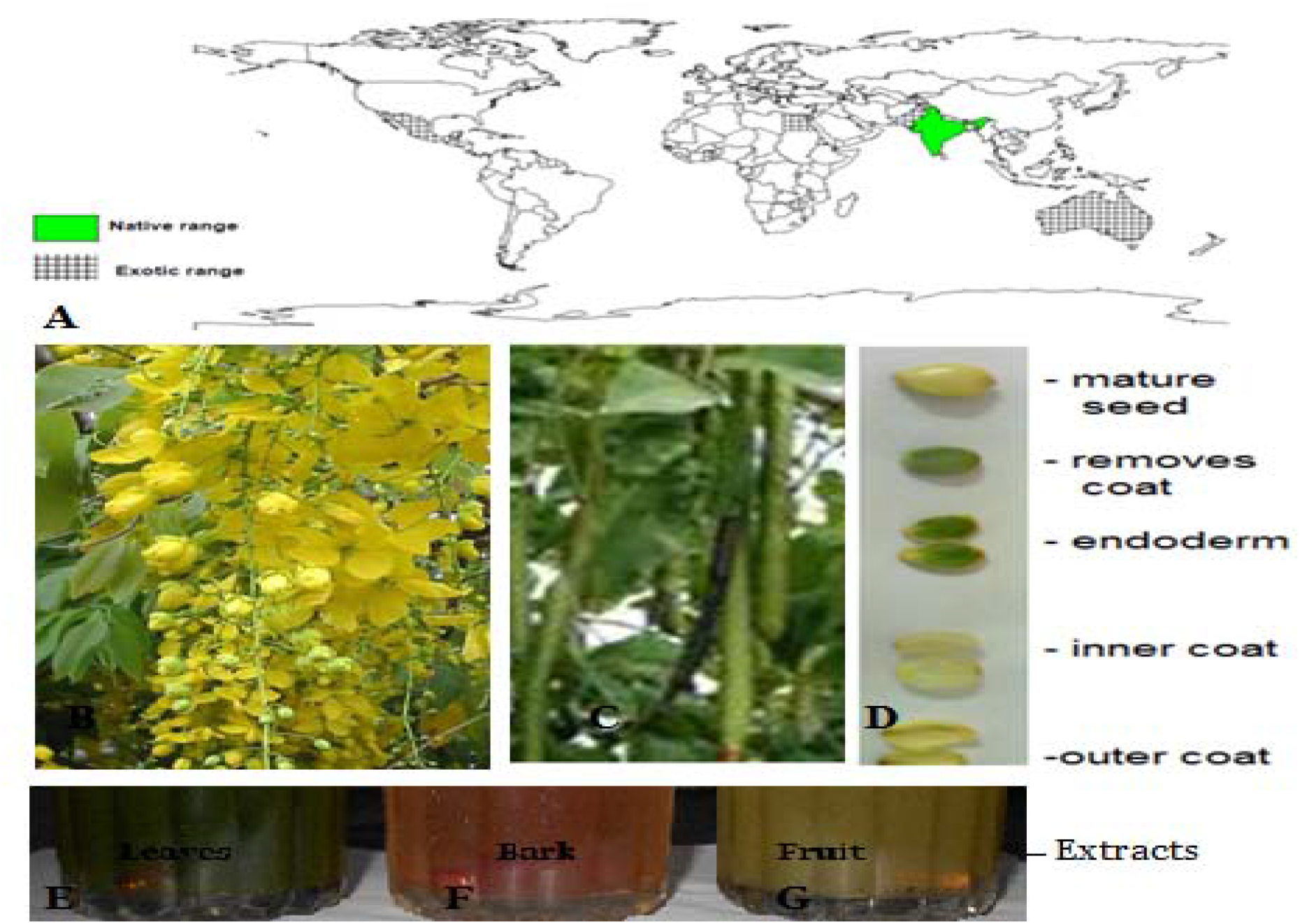
*Cassia fistula* Tree parts and extracts.

**Fig.2.**
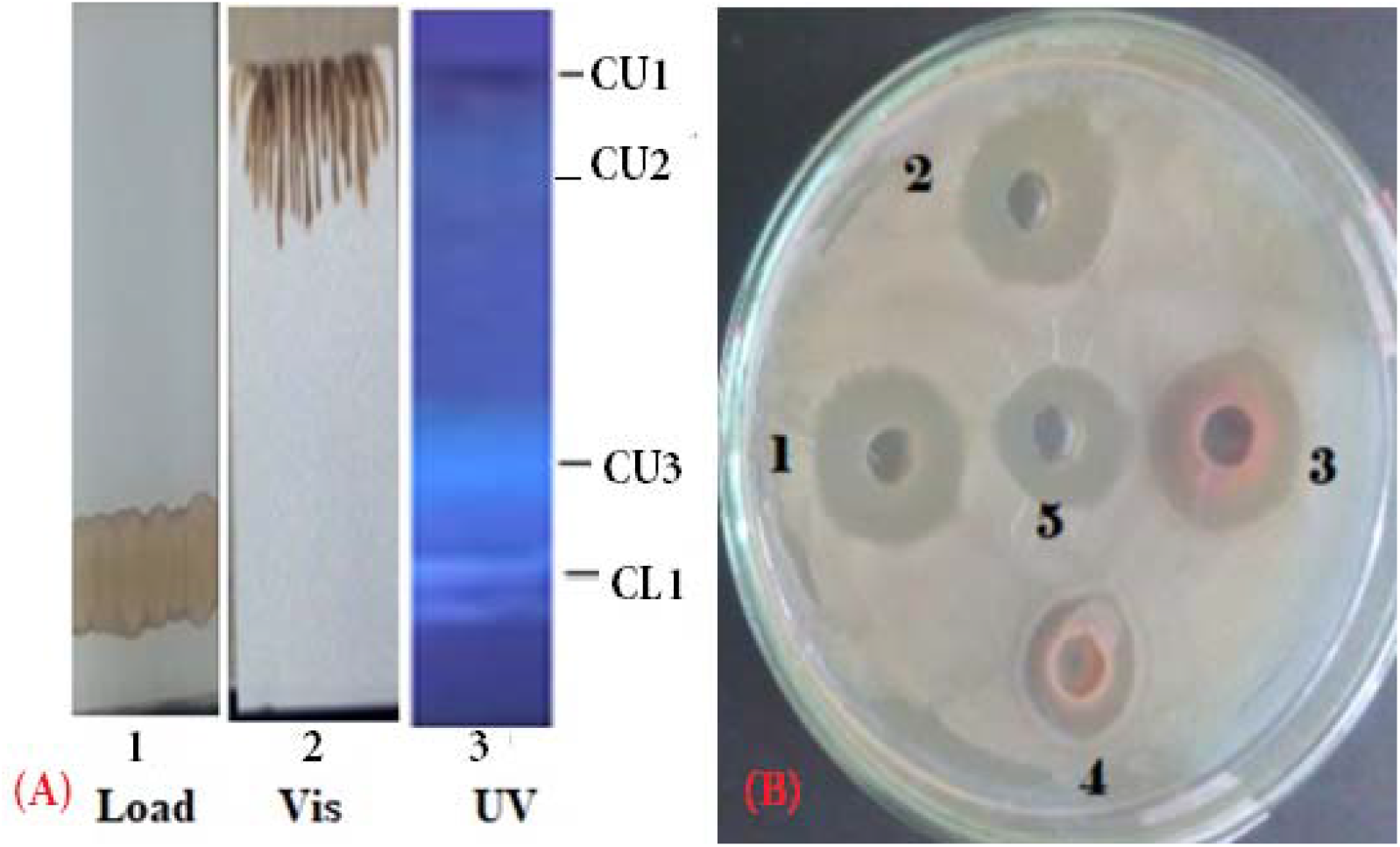
(A) Preparative Thin Layer Chromatography showing CU1 as gray fast moving major band. 1/6 of the TLC was shown here. CU3 also has anti-bacterial activity. (B) Kirby-Bauer Agar Hole Assay of TLC-purified different phyto-chemicals: 1. NU3; 2. VU3; 3. CU1; 4. CU1-HPLC purified; 5. Meropenem 20μl (1mg/ml)

### Purification of CU1 phyto-chemical

We purified the major phyto-chemicals CU1 and CU3 from 10-12 ml soluble ethanol extract using preparative TLC (20×15cm 8 plates and four solvent chambers) and 12 times such preparations were done and further TLC purification were repeated to make pure drug. Usually, we recovered ~300mg CU1/TLC purification scheme (figure-2A). We performed such purification scheme 3 times (in five years span) to perform all experiments (~1.2g CU1). CU1 chemical purified on HPLC C_18_ column at 3min and a minor contaminant at 6min (figure-3). Very low retention time of CU1 demonstrated a big molecule with hydrophobic nature like triterpenes.

**Fig. 3.**
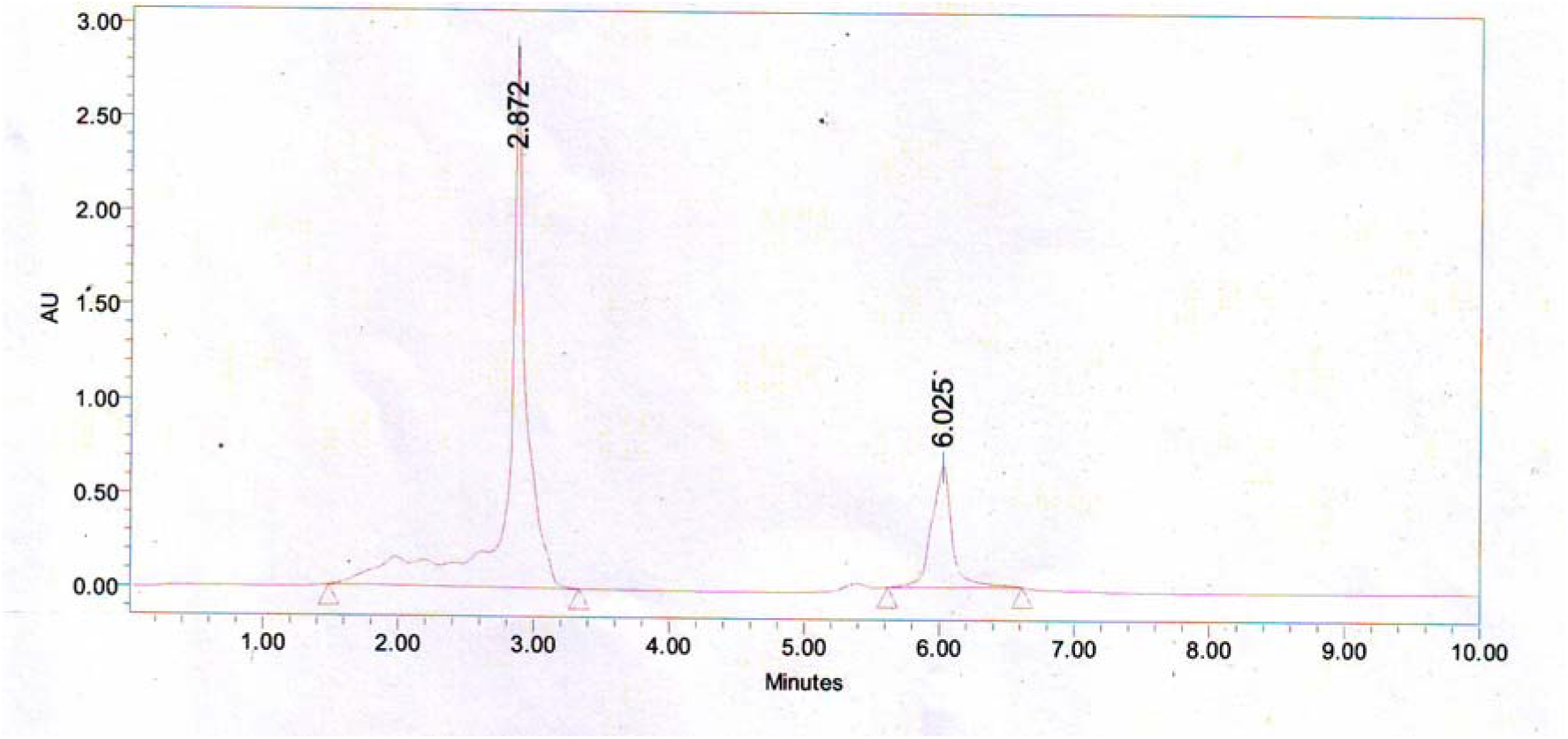
HPLC C_18_ column chromatography showing minor contaminant in TLC preparation and very fast elution time of CU1.

### Characterization of CU1 phyto-chemical and structure prediction

Chemical analysis also showed it had saponin character as well as triterpenes (figure-4). CHN elementary analysis gave 35.9% carbon and 5.5% hydrogen but no nitrogen indicating a saponin or glycosides. We confused for elementary data as 58.6% oxygen was impossible and we suspected a halogen derivative with at least 6 bromine atoms giving ~33.6% Br and ~25.2% Oxygen could be possible. The mass of CU1 was predicted 987.7173 daltons and from Mass spectra data (figure-5) we concluded that DBr 82MW deviations five derivatives (925.0639, 843.0582, 761.0507, 679.0442, 597.0367, 515.0299, 433.0221 daltons) indicating bromine atoms attached to carbon or -OH of polyphenol. CU1 Mass fragment of 325 may be triterpene plus a mono bromo-polyphenol and 987 Dalton mass was calculated as possible dimeric bromo-polyphenol derivative and a 79.5dalton major band may reflect Br+ ion but not shown here (figure-5). VU-Vis spectra gave 275nm peak with minor hinge at 578nm indicating a fused ring with polyphenolic substituent (figure-6). This interpretation was considered as phenolic compound and bromo-benzene both have primary peak at 210nm and secondary peaks at 270nm and 261nm respectively. So high peak at 210nm with secondary peak 275nm was justified for polyphenolic compound with multiple bromine where as for turpentine moiety further peak was shifted below 200nm was observed.

**Fig.4.**
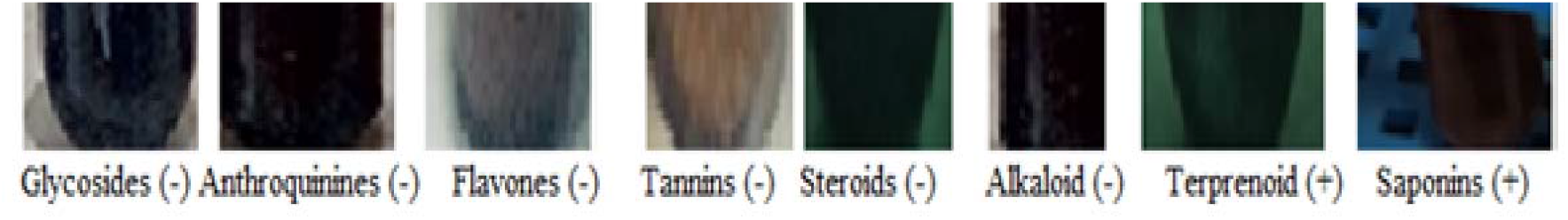
Biochemical assays of 1xTLC CU1 phyto-chemcal showing +ve for terpenetene and saponin.

**Fig.5.**
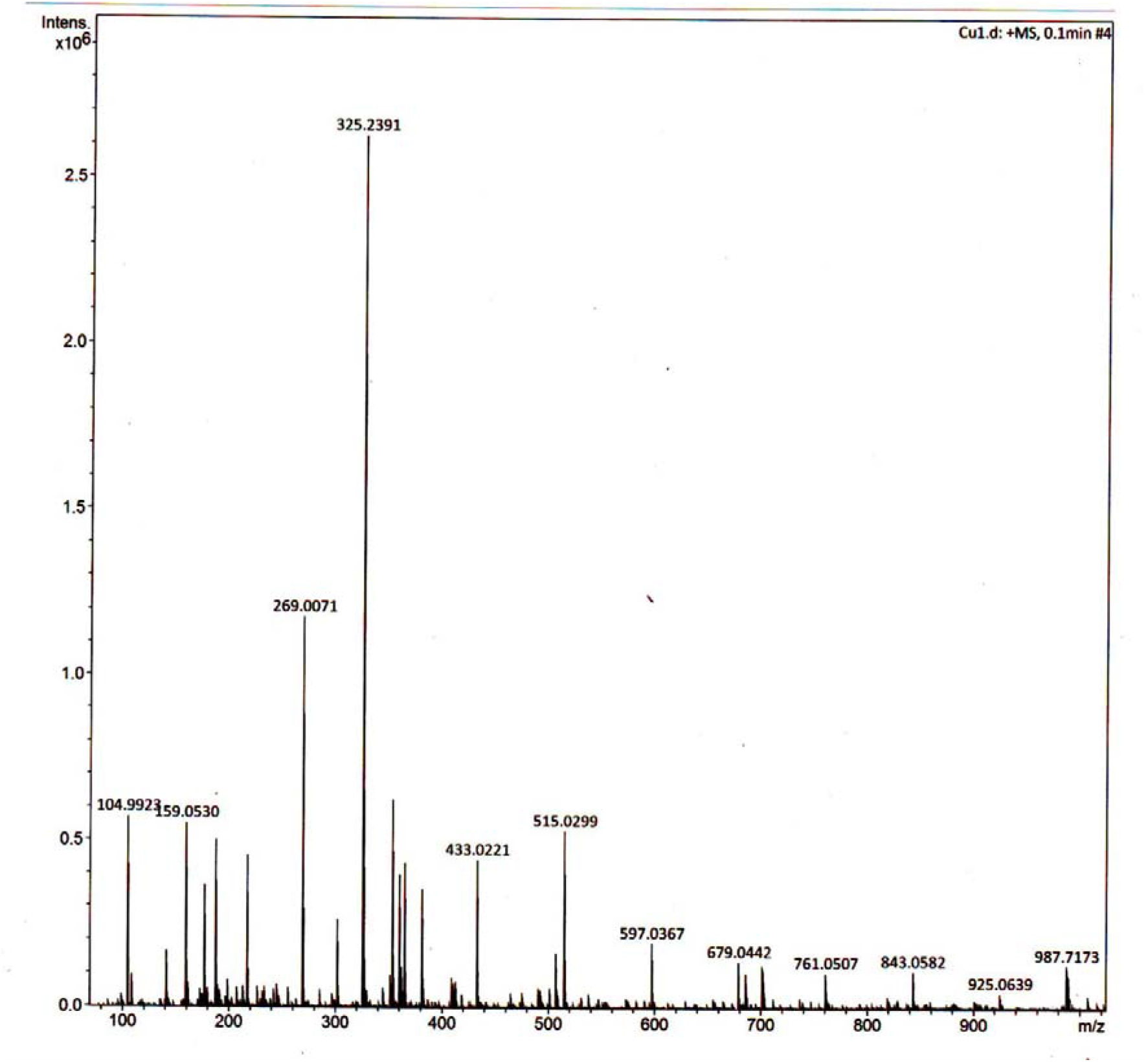
Mass spectra of CU1 phyto-chemical showing 987daltons MW. In this method, 100-1000MWmass limit but DBr band was detected at other method showing 79.5 Daltons major fragment.

**Fig.6.**
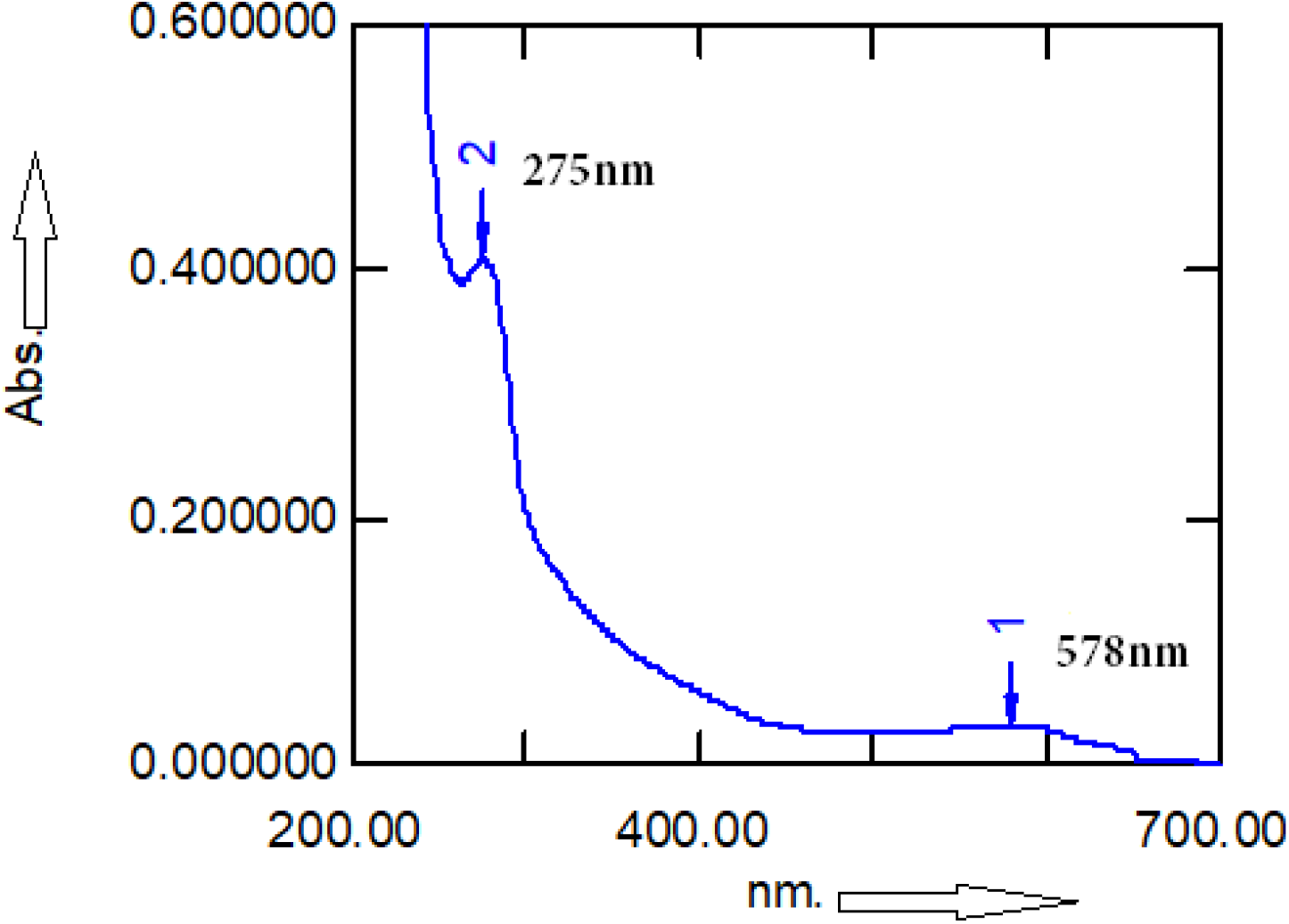
UV-Vis spectroscopy of TLC-purified CU1 showing peaks at 578nm and 275nm.

FT-IR suggested broad band at 3650-2980cm^−1^ for −OH (stretching, ~3500cm^−1^) and C-H (stretching, ~3000cm^−1^) where as two strong peaks at 1552cm^−1^ for aromatic C=C (stretching) and 1408cm^−1^ for phenol (-OH bending). Minor peaks at 2360cm^−1^ with 2340cm^−1^ shoulder represented terpenetine double bond. Similarly, medium peaks at 1246cm^−1^, 1159cm-1, 1105cm^−1^, ~837cm^−1^ and 651cm^−1^ were detected (figure-7). The absorptions peaks at 1246cm^−^ 1, 1159cm^−1^ and 1105cm^−1^ may be for secondary alcohol (-OH bending), at 837cm^−1^ likely aromatic hydrogen and at 651cm^−1^ with 619cm^−1^ shoulder likely represented for different C-Br. However, we did not prepare any derivative yet to confirm structure.

**Fig.7.**
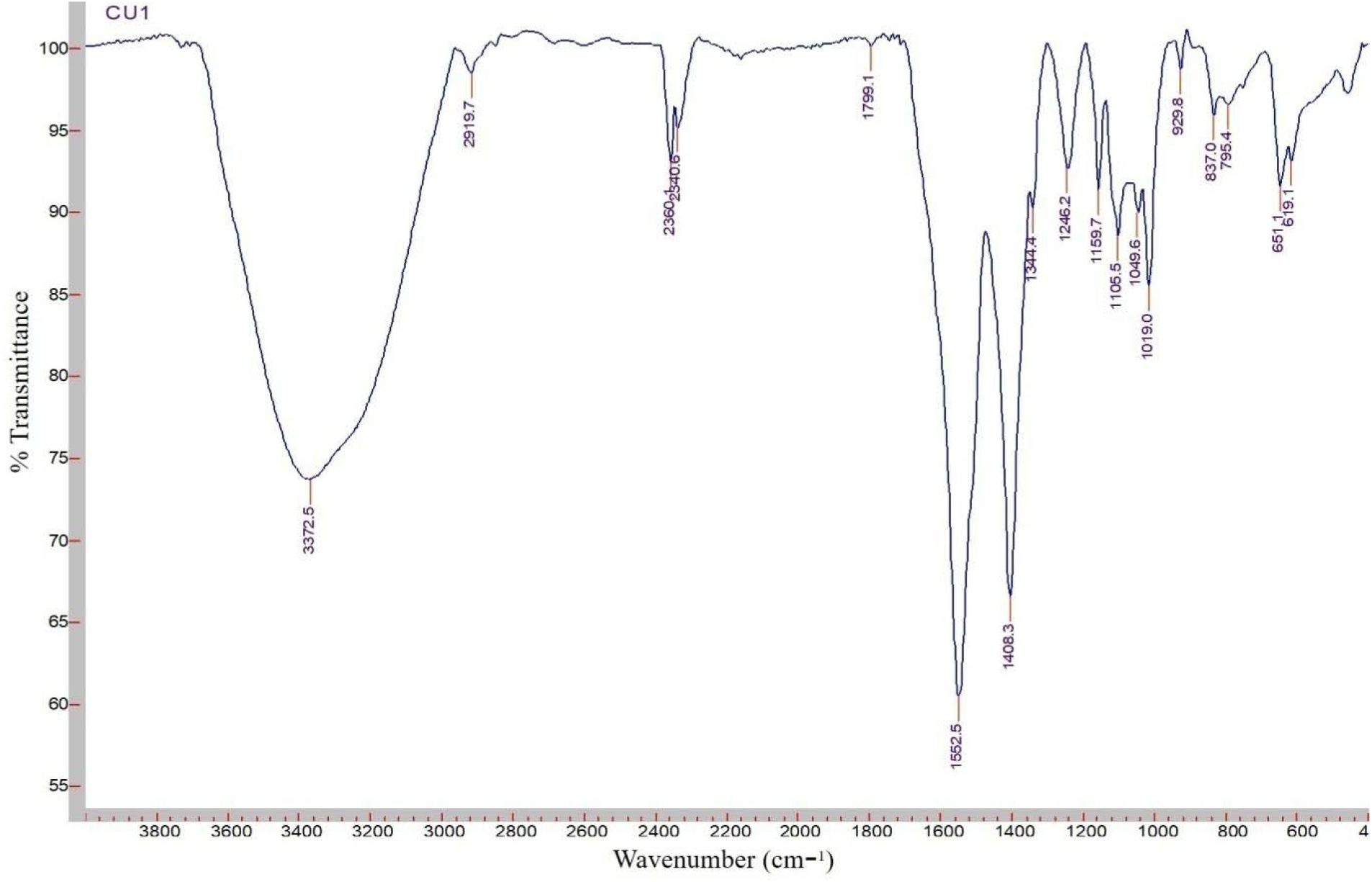
FT-IR spectra of CU1 phyto-chemical. A broad peak at 3000-3500cm-1 for −OH and strong peaks at 1552cm-1 and 1408cm-1are important for phenolics.

Proton-NMR confirmed polymeric phenol at δ 4.86-4.91 ppm and tetratet at δ3.57-3.61 ppm with phenolic bromo-substituents (figure-9). Benzoid group confirmed at δ 8.41-8.53 ppm where as sharp peaks at δ 3.3 ppm and δ 1.88 ppm confirmed phenolic bromo substituents (figure-8). A strong band at δ 8.41-8.53 ppm was evident in D_2_O as well as CD_3_OD but in CDCl_3_ H^1^ NMR a strong peak at δ7.262 ppm and medium peak at δ 1.588 ppm were evident (data not shown). However, peak at δ 1.88 ppm may link a fused C==O in the triterpene and for -CH_2_ at δ 0.11ppm (figure-8). The δ3.29-3.33 ppm bands in CD_3_OD was sharp at δ3.41ppm in D_2_O and δ2.75 ppm in CDCl_3_ (data not shown). This difference may reflect differential solubilisation and polar effects as triterpene moity is poor soluble water that polyphenol moity. Carbon-NMR identified a strong peak at δ=23.7ppm for many C-Br and at δ=165 ppm for a polybenzoid compound whereas weak peaks for -CH2- and = C-carbons of triterpene were detected at δ = 54 and 65ppm (figure-9). More analyses were necessary for complete chemical structure prediction (in process).

**Fig.8.**
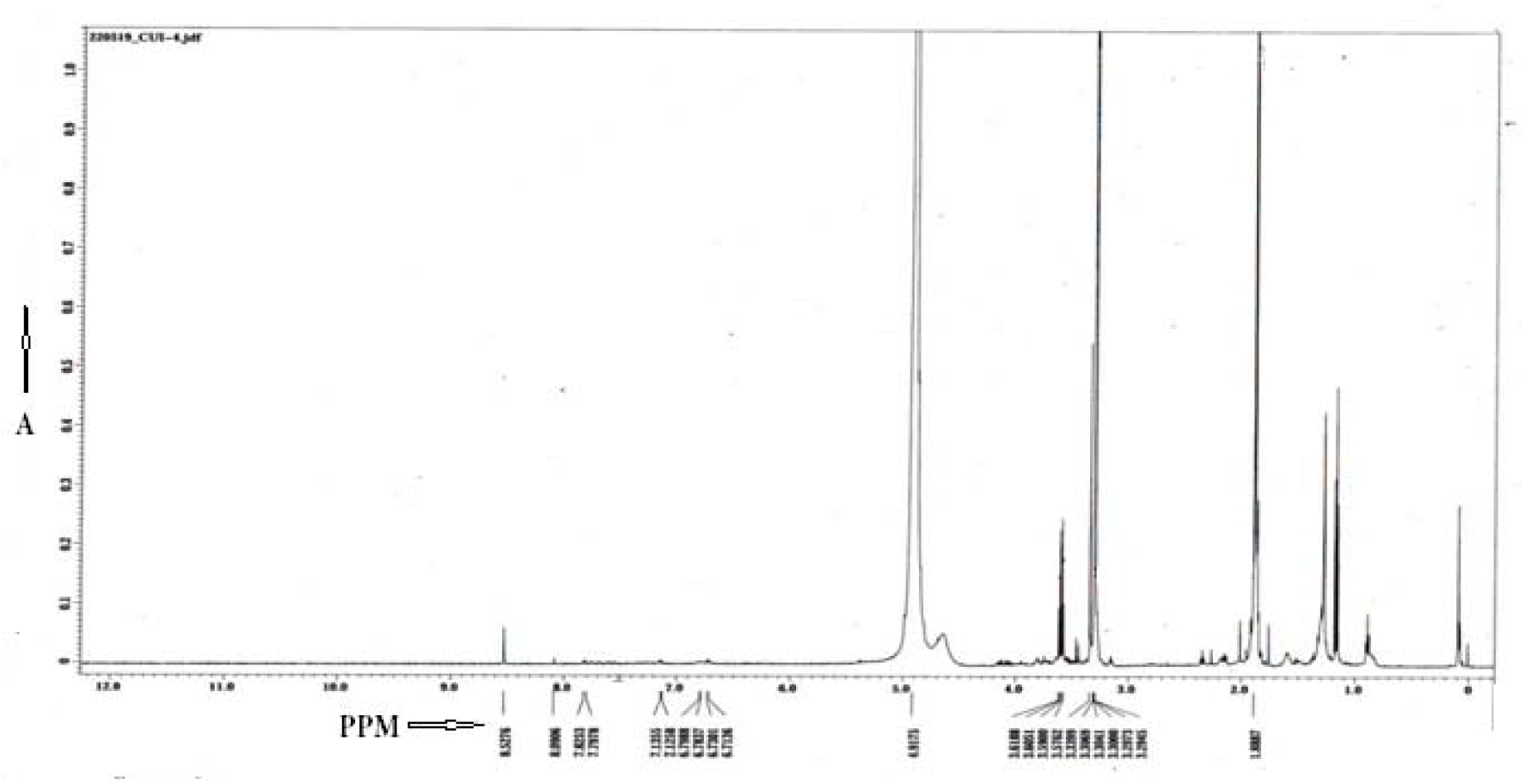
Proton-NMR spectra of CU1 phytochemical from Cassia fistula bark. Peaks at δ=8.4 ppm for polybenzoid and at δ= 4.9 ppm for polyphenolic -OH with δ=3.55 ppm for C-Br whereas at δ=1.88 for C=O absorption in triterpene.

**Fig.9.**
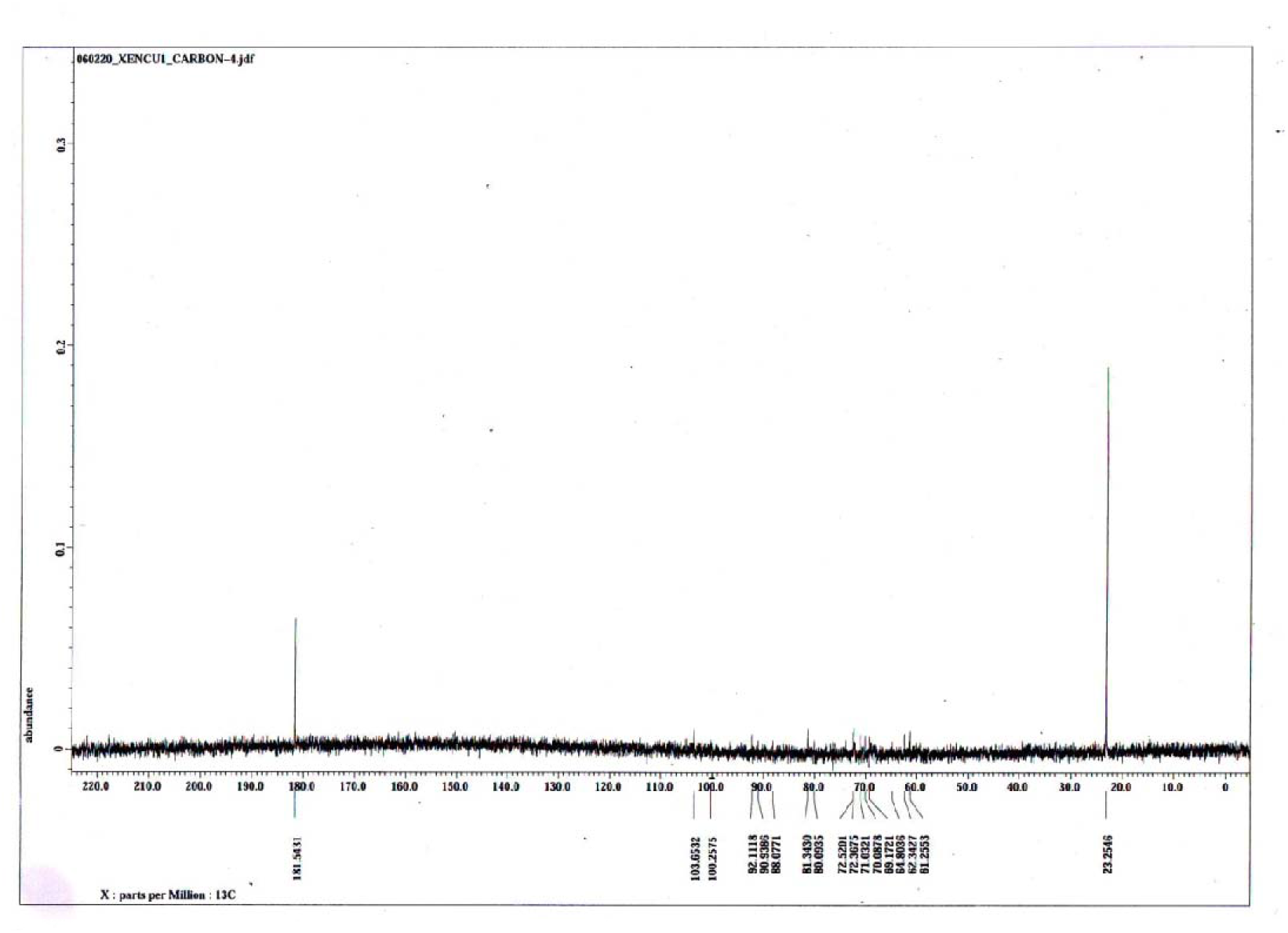
Carbon-NMR spectra of CU1 phyto-chemical. Strong peaks at δ=23.7ppm for many C-Br and δ=165ppm for polybenzoid compound are important. Small peaks for -CH2- and = C-carbons of triterpene were detected at δ = 54-65ppm.

### Transcription assays of 11 phyto-chemicals using E. coli RNA polymerase

A multiple-round non-specific transcription assay was performed using calf-thymus DNA as template, 2μci ^3^H-UTP as tracer and *E. coli* RNA polymerase. We found RNA polymerase activity was completely inhibited (complete-1325cpm, +Rif-63cpm, +CU1-75cpm, - Template DNA-52cpm) only with CU1 and NU2 phyto-chemicals but not with CU3, NU3, VU2, VU3, DU2, LU2, LU3 etc. The inhibitory activity of CU1 was comparable to drug refampicin and characterized further with priority (figure-10).

**Fig.10.**
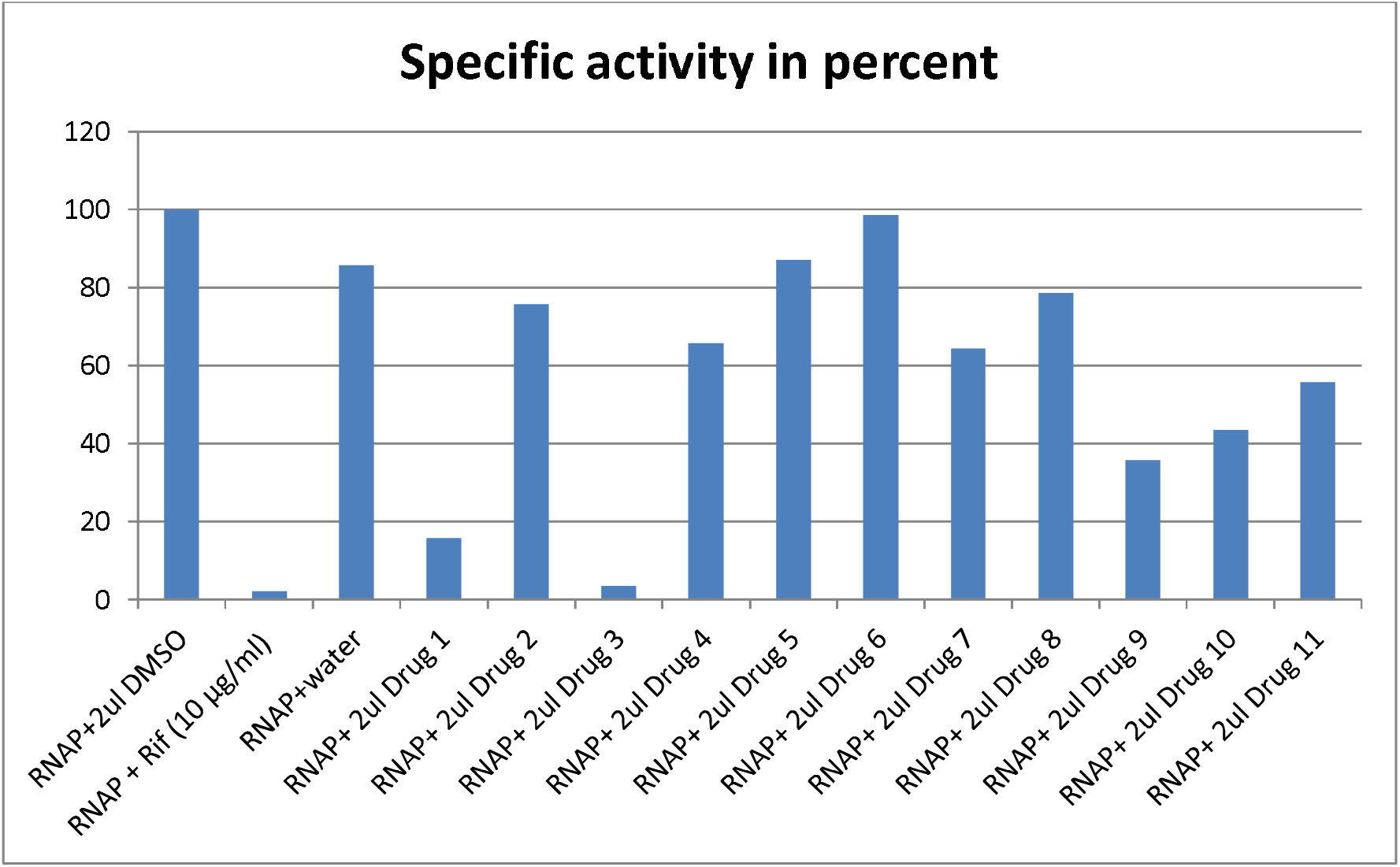
RNA polymerase assays of eleven bioactive phyto-chemicals isolated in our laboratory and discovery of target for CU1 (drug 3) from *Cassia fistula* bark and NU2 (drug 1) from *Suregada multiflora* root. Phytochemical concentrations are ~10mg/ml and standard is refampicin(Rif).

### Effect of CU1 on RNA polymerase of M. tuberculosis and E. coli

To test the effect of CU1 on transcription by RNAP of *Mycobacterium tuberculosis* a fluorescence based in vitro transcription assay was performed. Here, the RNA product formed after transcriptional elongation was bound to RiboGreen dye (Invitrogen, Carlsbad, CA), an ultrasensitive fluorescent nucleic acid stain used for quantitating RNA in solution. The fluorescence intensity in each reaction was then measured using a spectrofluorometer (Photon Technology International, HORIBA Scientific, Edison, NJ) at excitation and emission wavelengths of 500 and 525 nm, respectively. The results show that CU1 represses the transcriptional activity of *Mtb* RNAP*σ^A^*holo as well as *E. coli* RNA polymerase where refampicin is 2-3 time more active than CU1 (figure-11).

**Fig.11.**
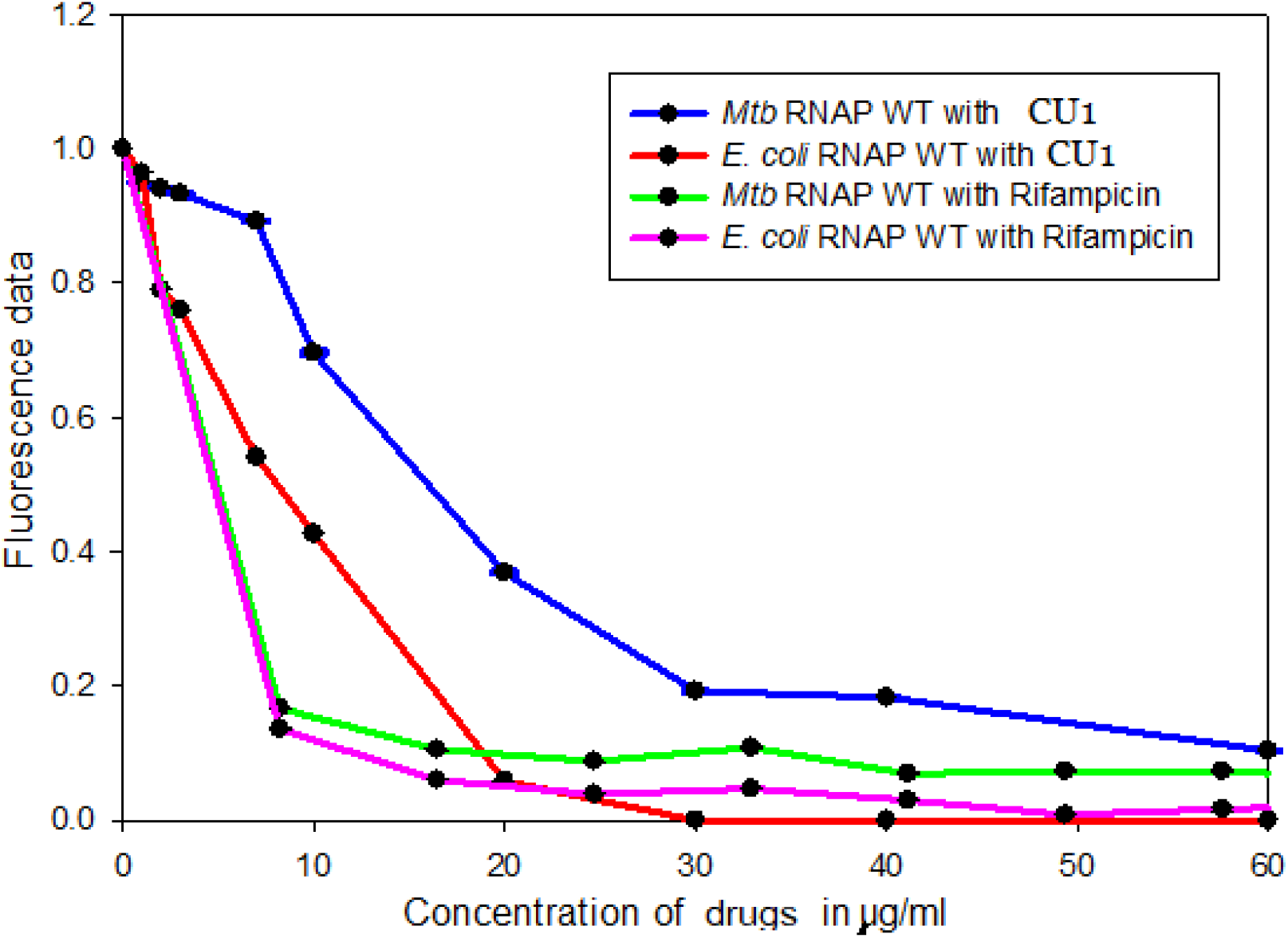
Kinetics of CU1 inhibition of *E. coli* and *M. tuberculosis* RNA polymerases as compared to refampicin drug.

To further test the transcriptional repression activity of CU1, radioactivity based in vitro transcription assay was performed using α^32p^-UTP. The *sinP3* promoter containing DNA used here generated a run-off transcript of length 70 nucleotides. The RNA products formed, incorporated the radio-labelled NTP present in the NTP mix (α^32p^-UTP) which helped to visualize the products after resolving in a 12% urea-PAGE. The results showed a repression of transcription with an increasing concentration of CU1 (figure-12 and figure-13). The estimated IC_50_ value of CU1 was found to be ~ 34μg/ml and in reactions containing 80 and 160 μg/ml CU1 the production of radio-labelled RNA was almost completely quenched (figure-13). The results obtained from these experiments confirmed that CU1 was a strong transcriptional repressor with *Mtb* RNAP. This is very interesting as MDR-TB is very much active in the Indian sub-continents.

**Fig.12.**
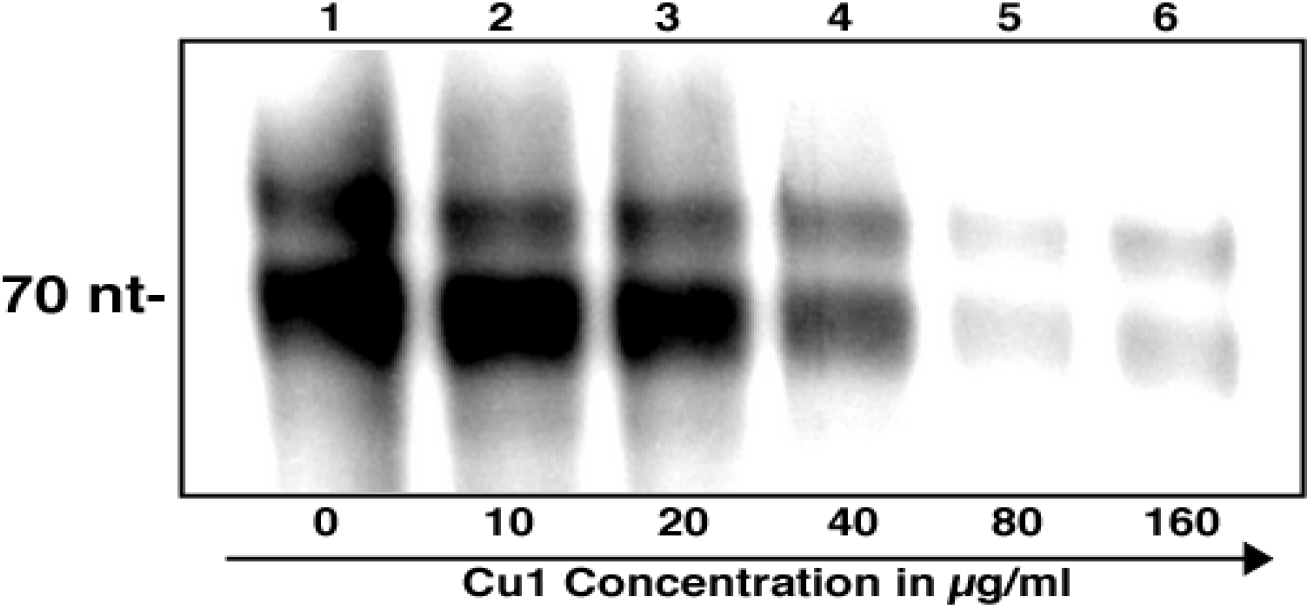
Radioactivity based in vitro transcription assay: 300nM *Mtb.*RNAP *σ^A^*holo was incubated with 1200nM *σ^A^*, 200nM *sinP3* promoter DNA, along with increasing concentrations of Cu1to form open promoter complex (RP_o_). Radiolabelled-NTP was added to initiate transcription. The estimated IC_50_ value of Cu1 was found to be ~ 34μg/ml. Concentrations of Cu1 used in the reactions (in μg/ml) are shown. Run-off transcript size is 70 nt.

**Fig.13.**
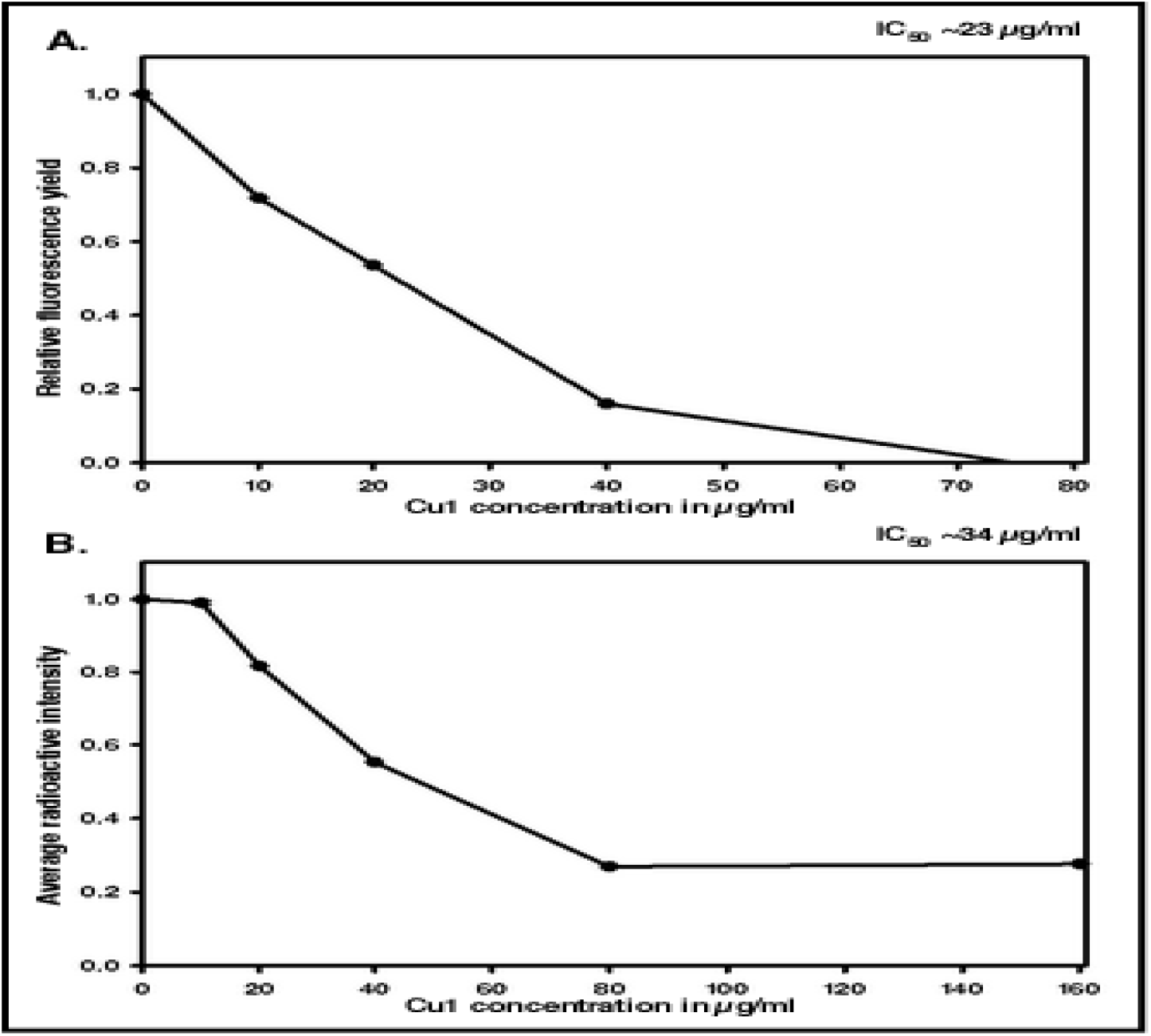
The effect of Cu1 on transcription by *Mtb* RNAP: **A.** Fluorescence based in vitro transcription assay: A plot of fluorescence intensity of RiboGreen bound to RNA in the presence of different concentrations of Cu1 (in μg/ml).The estimated IC_50_ value of Cu1 was found to be ~ 23μg/ml. The intensities were observed at excitation and emission wavelengths of 500 and 525 nm, respectively. **B.** Radioactivity based in vitro transcription assay. Average band intensity of transcripts in each lane from the Radioactivity based in vitro transcription assay was plotted for each concentration of CU1.

The next objective was to find out how CU1 was affecting the transcriptional process causing repression. For this study, Electrophoretic Mobility Shift Assay (EMSA) was done. For the initiation of transcription open promoter complex (RP_o_) formation is the first step, where the DNA forms a complex with RNAP. In this assay *Mtb* RNAP*σ^A^*holo was incubated with Cy5 labelled *sinP3* DNA to form RP_o_, in presence of increasing concentrations of the repressor CU1. The products were resolved in 5% native PAGE and visualized by fluorescence scanning. The results show that RPo formation reduced significantly with the increasing concentrations of CU1. The intensity of the shift denoting RP_o_ formation started to reduce in presence of 20 μg/ml of CU1 and eventually being completely absent in the presence of 160 μg/ml CU1 in the reaction (figure-14). This result showed that the transcriptional process was inhibited by CU1 at the initial open complex formation step of transcription, thereby causing transcriptional repression.

**Fig.14.**
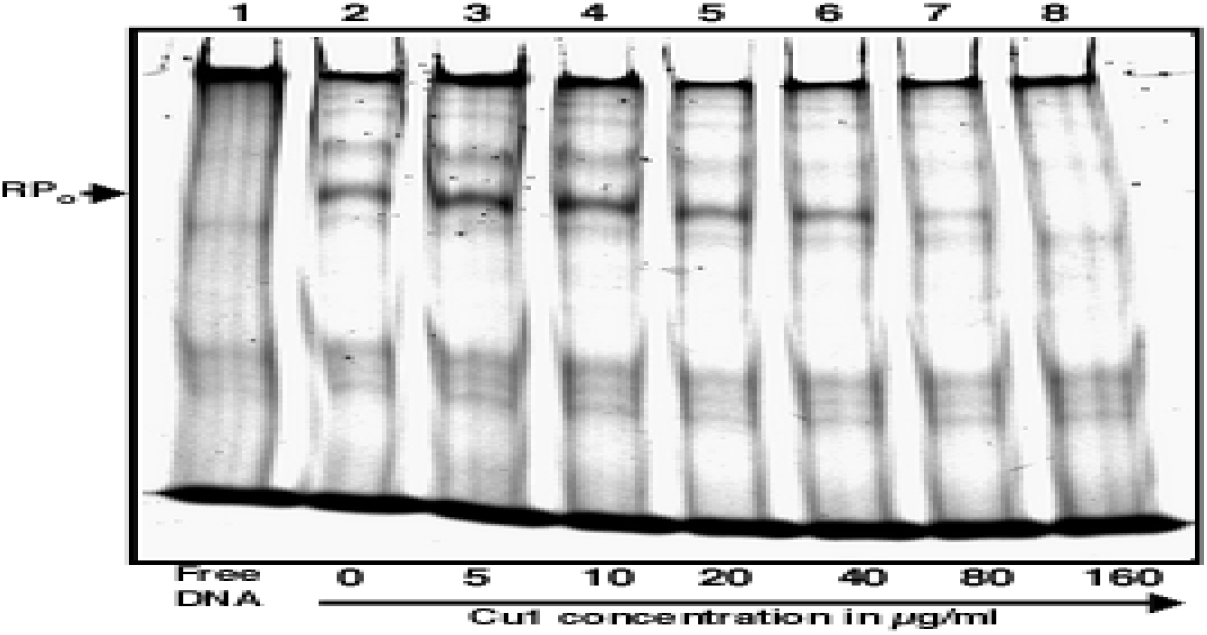
*Electrophoretic Mobility Shift Assay* (EMSA):. EMSA of the RPo with Cy5 labeled *sinP3* promoter DNA fragment in the presence of increasing concentrations of Cu1. For RP_o_ formation,1200nM*Mtb.* RNAP *σ^A^*holo was incubated with 400nM Cy5 labelled *sinP3* DNA and heparin (0.25mg/ml) along with increasing concentrations of the repressor Cu1 in subsequent lanes. Products were resolved by running in 5% native PAGE. Lane 1: Free DNA, Lanes 2-8: RP_o_ formation in presence of Cy5 labelled *sinP3* DNA and increasing concentrations of Cu1 (in μg/ml), up to 160 μg/ml.

### Effect of CU1 on DNA polymerase activity

The effectiveness of CU1 as a transcriptional inhibitor led to the investigation of its effect on bacterial DNA polymerase activity. For this study pUC19-*sinP3* single-stranded DNA template was annealed with Cy5 labelled complementary primer. Then, the primer was allowed to be extended by Klenow fragment (*E. coli* DNA polymerase; NEB) using the annealed single-stranded DNA as template, in presence of increasing concentrations of CU1. The products were then resolved in a 12% urea-PAGE for visualization. The results show no significant change in DNA replication products formed in the presence of increasing concentrations of CU1 (Figure-15). This shows that CU1 does not affect bacterial DNA polymerase activity. This also suggested that CU1 did not bind to DNA alone.

**Fig.15.**
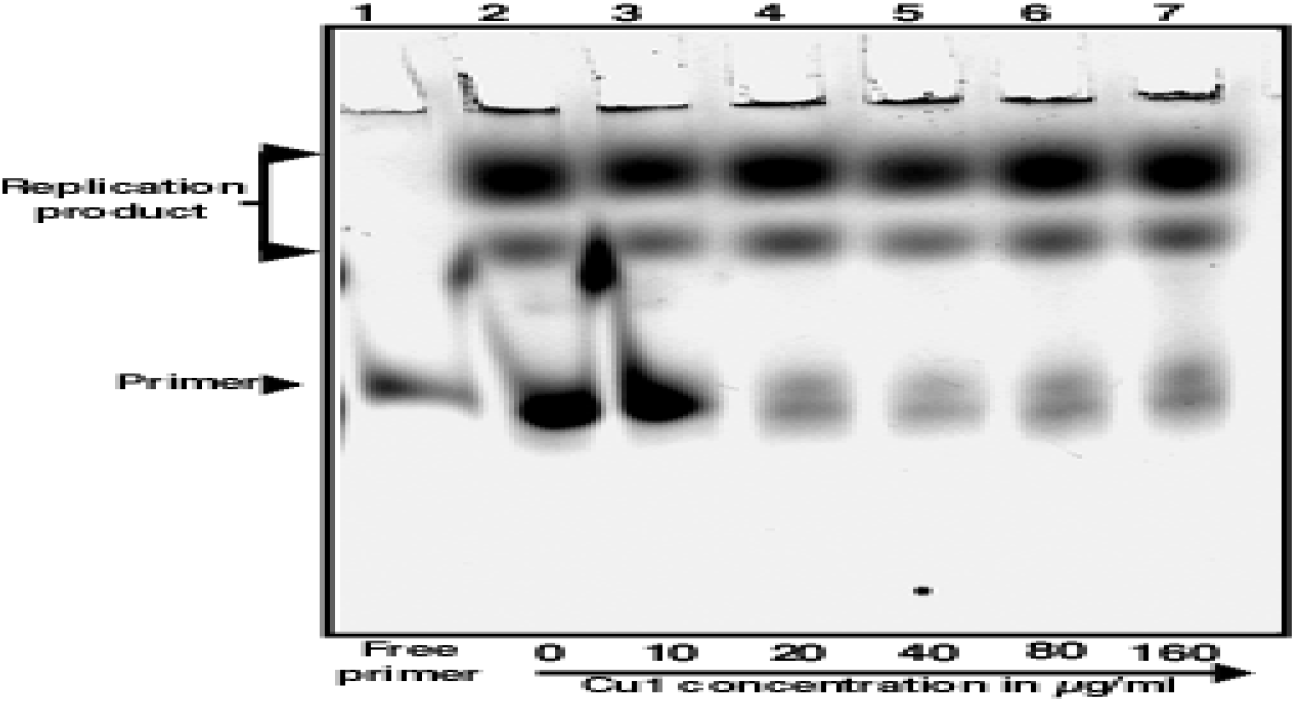
In vitro replication assay with the increasing concentration of Cu1. Single-stranded DNA template *pUC19-*sinP3 and its complementary Cy5 labelled primer were annealed and then incubated with increasing concentration of the repressor Cu1(in μg/ml). DNA polymerase, Klenow fragment (NEB) and 0.25mM dNTP mixture were then added to the reaction mixtures to initiate DNA polymerization. Lane 1: Free primer, Lanes 2-7: Replication product formation by Klenow fragment (NEB) in the presence of an increasing concentration of Cu1(in μg/ml).

## Discussion

Ancient drug development programmes were based on phyto-extracts. But after the discovery of multiple antibiotics in 1930s and their commercialization in 1940s pushed back auyrvedic drugs development. Penicillins, tetracyclines, fluoroquinolones and aminoglycosides drugs are easy to take, low cost and pure. Antibiotics saved mankind from cholera, typhoid, TB and other fungal and parasitic diseases between 1930-1980. Afterwards, we needed new derivatives of the drugs as penicillinases, oxacillinases, acetylases, phosphotransferases and adenyltransferases multidrug-resistant proteins appeared in plasmids as well as many drug efflux proteins like TetA/B/C, AcrAB, MexAB/CD/EF and MacAB. Multi-drug resistance has created a horror and phyto-drugs discovery has started again. Research indicated quinine, artiminisin, taxol, etoposide were good phyto-chemicals eliminating drug resistant malaria and cancer. This indicated a popular come back of phyto-drugs which are non-toxic and cheaper than synthetic antibiotics. We searched 80 plants to get new phyto-chemicals against multi-drug resistant bacteria which created a horror in society due to drug void. We pinpointed in previous papers that *Suregada multiflora* and *Cassia fistula* organic extracts were potential source of anti-bacterial phytochemicals [8, 12, 25]. In this communication, we proved that one abundant *Cassia fistula* photo-chemical (CU1) has RNA Polymerase target and we substantially gave UV-Vis, MASS, NMR and FTIR evidences for structure prediction of the photo-chemical as spoonin or terpentine linked to bromo-polyphenols. Due to corona virus pandemic our laboratory was closed for six months hindering our project. MDR-TB is increasing in India and drug resistant mechanism in Mycobacterium tuberculosis is complex. We have showed that *Mycobacterium* RNA polymerase was the target of CU1 with IC_50_ 34μg/ml. As the CU1 also inhibits *E. coli* RNA polymerase, other infectious Enterobacteriaceae like K*lebsiella. pneumoniae, Pseudomonas aerogenosa* and *Vibrio choleare* etc would be inhibited because *mdr* genes profiles in plasmids were indicated very similar and very similar RNA polymerase [36, 37]. Interestingly, *Mycobacterium tuberculosis* plasmids were rare and TB-specific drugs were different supporting authentic RNA polymerase inhibition by CU1 phyto-chemical led to a new drug discovery against TB [38]. Similarly, nature of *Neiserria gonorrhoeae* plasmids were different and many conventional drugs were reported non-functional indicating we needed more work on different in vivo assays in presence of CU1. CU2 or CU3 was not a inhibitor of RNA polymerase and at this point we did not predicts its molecular target.

We coined the name “MDR-CURE” which was a mixture of five medicinal plants extracts with traditional anti-oxidant and anti-inflamitory principals from 50% ethanolic solution of Neem bark and Haldi ryzome [8]. When we injected MDR bacteria into rats, ampicillin or cefotaxime failed to clear infection but MDR-Cure cleared the 80-90% bacterial load in blood within two days [8]. Our method of CU1 isolation is cheap and any one can make 95% pure phyto-chemical for human use by simple TLC method. In truth, heterogeneous phyto-chemicals will be a new treatment method to conquer MDR pathogenesis and the herbal drugs are non-toxic. Due to multi-resistance and worldwide contamination of MDR bacteria in water and air, WHO predicted a GDP loss of 3% worldwide in the coming decade. In such scenario, Corona infections have created panic worldwide claiming 40 million infections and >600000 deaths. In India, peoples are dying in fever, jaundice, diabetes, asthma and cancer due to lack of treatment during corona virus pandemic and drug void. We suspect some fever may be due to MDR-infections and US survey suggested that many corona virus treatment regime needed invasive antibiotics. In India drug sensitivity test was not mandatory for antibiotic prescription and old antibiotics were not curing diseases. Further, invasive 5^th^ generation antibiotics are costly and needs hospitalization. Such a pandemic is rare and if we use herbal medicine, many lives may be saved in poor and developing countries of Asia, Africa and Latin America.

We are engaged in determination of chemical structure of active phyto-chemical like CU1 of *Cassia fistula* tree. CU1 is water soluble and likely be a useful drug. Although in vivo assay demonstrated bacteriostatic nature of CU1 against *E. coli* in Luria-Burtoni medium (data not shown), However, CU1 plus CU3/NU3/VU3 combinations (MDR-Cure) were best for complete inhibition of MDR-Bacteria in LB medium similar to refampicin and other antibiotics. No apparent toxicity was found on rats and Molly fishes (data not shown) at 1% of phyto-extract or 10μg/ml. MDR-infections as well as MDR-TB and MDR-Typhoid have soared recently in India as well as other countries due to antibiotics failure. Thus, we claim heterogeneous phyto-antibiotics may be resourceful and safer as drug against MDR-bacteria. Multiple phyto-chemicals may also help to overcome the action of Tet, Acr, Mac and Mex types drug efflux proteins which also were detected in Kolkata MDR bacteria [8,26]. Twenty types of Beta-lactamases were isolated in large MDR conjugative plasmids which contained 10-15 *mdr* genes, few drug efflux genes and mutated sul1/2/3, PBPs and gyrAB proteins. Chloramphenicol acetyltransferase and related enzymes (AacC1 and AacA1) were mutated to twenty types drug acetyltransferases inactivating many aminoglycosides (kanamycin) and fluoroquinolones (ciprofloxacin). Similarly, early detected streptomycin-phosphotransferase (StrA and StrB) were mutated to fifteen types APH phosphotransferases inactivating kanamycin, amikacin and gentamycin [36–39]. Further, specific drug-inactivating mechanism also creating problem of TB therapy and Gonorrhoea control where as Cholera, Pneumonia and Typhoid were mostly drug resistant and 4-8 antibiotics were needed to clear infections [40]. Repeated drugs use may damage liver and kidney and may be reason for other metabolic diseases like diabetes and hypertension. So, herbal therapy is must and research must be augmented in that area. However, India has started plantation in large areas (Herbal Garden). Phage therapy may also be welcome but phage resistant factor is a big obstacle. We found Kolkata drain has some bacteriophages but hard to get phage particles in large number due to resistant factor during large scale re-infection assay (data not shown). Genetic engineering applications are used to make recombinant phages to overcome resistant factor but most data are hidden whereas virus mutagenesis is not a safe method. Genetic drugs (antisense, ribozyme) and gene therapy technologies will be costly and poor peoples of Asia, Africa and Latin America and poor peoples have limited access to such costly therapy (41). Hundred thousand peoples were dragged into nasty poverty line every year due to high cost of antibiotics, anti-cancer and psychoactive drugs. In that scenario, MDR-Cure must be given priority by WHO and Leaders of G-20 Nations [42].

## Conclusion

Repeated oral antibiotics uses have been reported worldwide which drives more *mdr* gene creation and drug void. Excessive synthetic drugs also damage the liver and kidney increasing metabolic diseases like high pressure, diabetes, hypertension and cancer. Thus, the need of MDR-Cure herbal drug against MDR pathogenesis is necessary and timing. We preliminary have characterized *Cassia fistula* (golden shower) CU1 phyto-chemical as complex structure of turpentine linked to bromopolyphenol but complete structure yet to determine. We suspected that 2-3 bromine might be also attached to triterpine moiety. How bromopolyphenol moiety was linked to triterpine was not sure at this point! As we also suspected a “S” atom (-SO_3_H group) in CU1 molecule which complicated our structure prediction. We are in process of characterizing other phyto-chemicals from *Suregada multiflora*, *Jatropha gossipiifolia* and *Shorea robusta* etc (figure-16). Such bark is available in large amount from a medium sized tree which can grow in unfertile land and no water treatment was found necessary during summer. It was reported that many pure phyto-chemicals are more toxic to animal cells that crude extract. In truth herbal drugs means crude drugs and licence for human use could be obtained easily. However, taxol, artiminisin, quinine types phyto-drugs have used as 95% pure similar to antibiotics (11). Thus, our approached to purify and characterize the CU1 phyto-chemical in detail to formulate as MDR-Cure drug is justified. Pure TLC purified CU1 was very active and such drug could be obtained at low cost. We have selected six plants and spices to make MDR-Cure and such drugs in combination will be effective against pan drug resistant bacteria. Definitely more work necessary to pinpoint final structure of CU1 and also clinical study necessary before its use as prescription drug. Our MDR-cure lotion cures many human MDR nail infection (unpublished data). Abundant CU1 antibiotic is novel, water soluble and will also help to understand the molecular mechanism of RNA synthesis in bacteria as there are very few drugs available today for RNA polymerase inhibition study and drug design against MDR bacteria.

**Fig.16.**
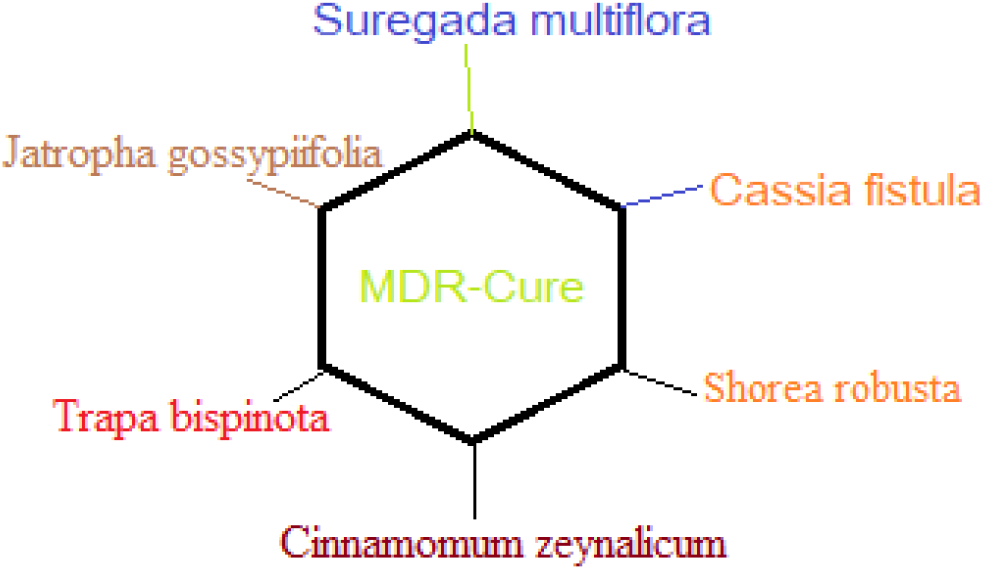
Heterogeneous phyto-chemicals in MDR-Cure and drug development from Indian medicinal plants against superbugs. *S. multiflora* (naranga) bark and root, *C. fistula* (bandor-lathi) bark, *J. gossypifolia* (veranda) root, *T. bispinota* (panifal) fruit peel, *S. robusta* (sal) inner bark and *C. zeynalicum* flower bud (derchini) were extracted in ethanol, concentrated five times before Kirby-Bauer agar-hole anti-bacterial assay. We are sure that any multi-drug resistant infection will be cured by such extracts in combination where as purified phyto-chemicals using preparative TLC are equally good.

## Acknowledgement

We are sad to state the sudden demise of Dr J. B. Medda, President of Oriental Association of Education and Research. AKC thanks Prof Dipankar Chatterjee of Indian Institute of Science, Bangalore for allowing his laboratory to test 11 phyto-chemicals on *E. coli* RNA polymerase. AKC thanks Dr Ram Maji of Indian Institute of Chemical Biology for HPLC and Prof Sujoy Dasgupta, Bose Institute for helping NMR spectroscopy. Funding from DST-SERB to Dr. JM and OAER Institutional funding to Dr AKC are acknowledged. Special thanks to Council of Scientific & Industrial Research, Govt of India for providing senior research fellowship to Surajit Saha.

## Ethical Issues

No human patient was used. Rat and Molly fish experiments with CU1 were discussed with Institutional Ethics Committee.

## Conflict of Interest

All authors declared no conflict of interest.

## Notes

### Competing Interest Statement

The authors have declared no competing interest.

